# Learning-accelerated Discovery of Immune-Tumour Interactions

**DOI:** 10.1101/573972

**Authors:** Jonathan Ozik, Nicholson Collier, Randy Heiland, Gary An, Paul Macklin

## Abstract

We present an integrated framework for enabling dynamic exploration of design spaces for cancer immunotherapies with detailed dynamical simulation models on high-performance computing resources. Our framework combines PhysiCell, an open source agent-based simulation platform for cancer and other multicellular systems, and EMEWS, an open source platform for extreme-scale model exploration. We build an agent-based model of immunosurveillance against heterogeneous tumours, which includes spatial dynamics of stochastic tumour-immune contact interactions. We implement active learning and genetic algorithms using high-performance computing workflows to adaptively sample the model parameter space and iteratively discover optimal cancer regression regions within biological and clinical constraints.

## Introduction

### The Translational Dilemma in Cancer

Immunotherapy—which retunes the body’s immune system to control cancer progression or eliminate it altogether—is one of the most promising cancer treatment strategies to emerge in the past ten years^1^. In 2010, Hodi et al. reported that some patients with metastatic melanoma had improved survival time after potentiating anti-tumour T-cell response (by targeting CTLA-4)^2^. Durable (and even complete) responses were observed in a significant fraction of those receiving the immune treatment^2^. More recent immunotherapies targeting PD-1 or PD-L1 have significantly improved 3-year overall survival and progression-free survival time in melanoma^3^. Similar advances have been reported in diverse cancers such as non small-cell lung carcinoma, hepatocellular carcinoma, and renal carcinoma^4,5^.

However, cancer immunotherapies do not benefit all patients equally: only 10-20% of patients receiving immunotherapy experience durable, partial or complete responses, and far more experience only temporary responses or none whatsoever^4,5^. Much work has focused on finding biomarkers to identify the 20% who can best benefit from immunotherapy^6^, while many others have focused on improving immunogenicity^4,5^.

This work has been complicated by the complex dynamics of tumour-immune interactions that change through the stages of immunoediting^7^. In the elimination phase, the innate and adoptive responses are effective in eliminating cancer cells as they emerge. In the equilibrium phase, some tumour cells escape, but the immune system keeps the overall tumour cell population under control. In the final escape phase, tumour cells evolve to evade immune recognition or even subvert immune processes (e.g., inflammation) to drive further growth^7,8^. Immune interactions can both harm and help growing tumours, thus complicating efforts to develop immunotherapy strategies. Moreover, even effective immunotherapeutic responses can follow an initial period of tumour growth or pseudoprogression^9^.

Dynamical mathematical models can pierce the complexity of tumour-immune interactions and inform our therapeutic strategies^10–12^. Due to the dynamical nature of individual immune cells, the nuances of individual immune-immune and immune-cancer cell interactions, and tumour cell heterogeneity, it is advantageous to use agent-based models (ABMs) to mathematically model individual cancer and immune cells (each with individual positions, states, and immune characteristics), rather than simulate populations of cells with blurred positions and properties^13^.

Numerous agent-based models have been developed to study cancer-immune interactions and immunotherapy. (See the excellent recent review by Norton et al. 2019^14^, and our own recent work by Ghaffarizadeh et al.^15^ and Ozik et al.^16^.) By adjusting model parameters and simulation rules, we can explore the characteristics of successful and unsuccessful treatments, and learn how the “best policies” vary with a patient’s tumour characteristics^16–18^.

If we can identify the “behavioural rules” of a successful immunotherapy, we can seek molecular interventions to implement those rules. For example, we could modify immune receptor binding efficiencies and protein lifetimes to influence downstream behaviours such as contact-based interactions^19–21^, or introduce virotherapies that introduce new genes to a system^22^. Immune cells could be engineered to secrete co-stimulatory ligands to modify differentiation, activation, cell trafficking, and other critical multicellular behaviours^23^. T-cell motility varies from random to strongly directed to help balance exploration and exploitation, based in part upon how well cells can detect gradients of chemotactic signals^24–26^. Chemotactic motility could potentially be modulated by modifying chemotactic receptor expression, receptor-ligand binding efficiency, or receptor trafficking characteristics. Cancer immunotherapy could thus benefit from simultaneously employing molecular approaches (What medicinal chemistry can be employed to target specific molecular biology?) and multicellular systems-level approaches (What immune cell rules lead to the best cancer control and remission?).

### Computational challenges: Scalability

However, the benefits that arise from ABMs being able to reproduce the complexity of their referent biological systems also present challenges to how they are used. Specifically, one of the longstanding benefits of mathematical models of biological processes is their ability to use power of abstraction to aid in the identification of fundamental principles governing these systems and provide access to the entire world of formal analysis to enhance understanding. Alternatively, ABMs, as well as other types of complex, dynamic, multi-scale models, are themselves highly complex objects that are, to a great degree, not reducible to a formally analytical form. As a result ABMs are generally treated as experimental objects, used in simulation experiments similar to their biological brethren, and their overall behaviour can only be evaluated by the execution of very large numbers of simulations, a multi-faceted process we refer to as model exploration (ME). ME is a near-ubiquitous component in the development of models and algorithms; as applied to ABMs and other multi-scale models, it often involves an iterative workflow where simulation experiments are carried out across a large range of parameter values for purposes such as model calibration, model optimization or model behaviour characterization. Model output after one set of simulation experiments are evaluated against some predetermined metric, which informs the next iteration of simulation experiments. Examples of ME algorithms include Active Learning (AL)^27^ and Genetic Algorithms (GA)^28^, both of which are used in ME studies in this paper. While advances in high-performance computing (HPC) can allow for the parallelization of certain aspects of this process, resulting in high-throughput dynamic knowledge representation and hypothesis evaluation to address a current bottleneck in the Scientific Cycle^29,30^, we propose that the ME process itself can be enhanced with a computational framework^31^. The remainder of this paper presents a ME workflow implemented via an integration between an existing toolkit for creating ABMs of cancer, PhysiCell, and an HPC ME workflow framework, Extreme-scale Model Exploration with Swift (EMEWS); the developed workflow investigates the characterization of the parameter space of an abstract model of immunotherapy on a generic tumour model to find optimal cancer regression regions.

## Description of Bio-ABMs and PhysiCell

### Agent-based modelling in cancer immunology

In agent-based models, each individual cell is a software agent: an autonomous object with individual parameter values, state, and methods (functions) to govern behaviour in a virtual environment. In the context of simulated cancer immunology, the agents are cancer and immune cells, their parameters describe biological properties like birth and death rates and immunogenicity, the state can include the cell’s position and cell cycle status, the virtual environment is the tissue microenvironment, and the methods can control entry into the cell cycle, cell motility, and other key biological processes. Introducing new cell types is a matter of defining their individual functions and setting their parameters; thus, modellers can readily build simulation models that can explore the emergent dynamics of various immune cell types as they interact with each other and cancer cells in 3-D tissue microenvironments. This affords the possibility of identifying key biophysical parameter constraints in improving cancer immunotherapies.

To date, most mathematical modelling of cancer-immune interactions have used non-spatial models (i.e., systems of differential equations), molecular-scale models of signaling dynamics, or lattice-based agent-based models that could not readily investigate the impact of mechanical interactions between tumour and immune cells. See Norton et al.^14^ for a review of agent-based simulation models of tumour immune microenvironments, and Metzcar et al.^13^ for a broader overview of cell-based computational modelling in cancer biology. In the work below, we focus on previously unexplored mechanical, spatial, and stochastic aspects of tumour-immune contact interactions.

### PhysiCell: a platform for multicellular systems biology

In Ghaffarizadeh et al.^15^, we developed PhysiCell, a general purpose simulation platform for multicellular systems biology. In this C++ modelling framework, each cell is an off-lattice agent with motion governed by the balance of adhesive, repulsive, motile, and drag-like forces. Each cell has an independent cell cycle state (including volume changes), can perform biased random migration with user-programmed functions, and can progress through apoptotic and necrotic death processes. Modellers can attach customized data and C++ functions to each agent, and dynamically modify these data and functions in individual cells throughout the duration of a simulation. This allows the framework to be very closely tailored to selected biological problems. Its modular design allows further customization by open source contributors. For example, Letort et al. recently integrated Boolean signalling networks into each cell agent to model molecular-scale processes^32^.

In most models, cell behaviour is linked to the values and gradients of diffusing substrates, such as oxygen-dependent cell cycle entry and necrosis and chemotaxis towards signalling factors. To facilitate this, PhysiCell uses BioFVM^33^ to solve 3-D diffusion equations for one or many diffusible factors (typically 1-10 factors), with automated integration with the cell agents. Each cell can readily secrete or uptake from the chemical microenvironment, or sample the value or gradient of any or all substrates.

PhysiCell is cross-platform compatible: models can be compiled and run on Linux, macOS, Windows, and other operating systems with little to no modification. The framework has been parallelized with OpenMP and tuned to run 3-D simulations of 10^5^ or more cells on desktop workstations.

### PhysiCell model of cancer-immune contact dynamics

In Ghaffarizadeh et al.^15^, we introduced a simple model of 3-D immunosurveillance against heterogeneous tumours, with a special focus on the spatial dynamics of stochastic tumour-immune contact interactions. In the model, each cancer cell has a mutant “oncoprotein” *p* which drives proliferation: the greater the expression of *p*, the more likely the cell cycles and divides. In the absence of other selective pressures, the cells with the greatest *p* expression clonally expand and dominate the dynamics of the tumour. Under the simplifying assumption that a highly-expressed mutant protein would be reflected as a more immunogenic peptide signature on major histocompatibility complexes (MHCs)^34^, we modelled each cell’s immunogenicity as proportional to *p*.

To model immunosurveillance, after simulating 14 days of growth we introduced generic immune cell agents that move towards tumour cells by chemotaxis (a random biased migration towards a cell-released chemical factor), test for contact with cells, stochastically form spring-like adhesions to any cell in close contact, and then test for immunogenicity. While adhered to a target cell, the immune cell agent attempts to induce apoptosis (e.g., by the FAS receptor pathway^35^) with a probability that scales linearly with immunogenicity. If successful, the tumour cell undergoes apoptosis, while the immune agent detaches and resumes its chemotactic search for additional tumour cell targets. If the immune cell does not kill the tumour cell, it remains attached while making further attempts to induce apoptosis until either succeeding or reaching a maximum attachment lifetime, after which it detaches without inducing apoptosis. See Ghaffarizadeh et al.^15^ (2018) for further technical and mathematical model details.

The model presented in this paper was implemented using PhysiCell Version 1.4.1, and modified the model from Ghaffarizadeh et al.^15^ and Ozik et al.^16^ to allow selection of 2-D or 3-D simulations. The full source code is available at GitHub; see the link in the Electronic Supplementary Information.

A 4K-resolution video of the 3-D model was published as part of Ghaffarizadeh et al. (2018), and can be viewed at https://www.youtube.com/watch?v=nJ2urSm4ilU. Readers can interactively run and explore the 2-D model at http://nanohub.org/tools/pc4cancerimmune, built using xml2jupyter^36†^.

In Ghaffarizadeh et al.^15^, we performed one large 3-D simulation of this model, finding that stochastic immune cell migration had a major impact on this system by increasing spatial mixing between tumour and immune cells, potentially contributing to more successful immune responses. However, to further understand this system would require hundreds or thousands of additional simulations.

### Preliminary model exploration

In Ozik et al.^16^, we explored a three-dimensional parameter space to further investigate the role of stochasticity in this model. The parameters investigated consisted of:

- **immune cell attachment rate:** the rate at which an immune cell can form an adhesive bond with another cell in close contact;
- **immune cell attachment lifetime:** the mean time the immune cell spends attached to a target cell before detaching and resuming its search for targets; and
- **migration bias** (with 0 ≤ *bias* ≤ 1). If the bias is 0, migration is purely Brownian, while a bias of 1 indicates deterministic chemotaxis. Intermediate values give a biased chemotactic random walk.

We discretized the parameter space into 27 parameter sets (low, medium, and high values for each parameter), with multiple simulation replicates (with different random seeds) per parameter set. We found that both the attachment rate and attachment lifetimes had threshold effects: once the parameters were sufficiently high, further increases did not significantly improve the immune response. However, the bias parameter was markedly non-monotonic: either decreasing bias (leading to more *exploration* by immune cells by more random tumour-immune mixing) or increasing the bias parameter (leading to more *exploitation* by immune cells by directly moving to the closest tumour cells) led to an improved immune response compared to our original simulation^15^.

However, this work was only a first step: it did not further explore the parameter space to find phase transitions in model behaviour or to find optimal parameter sets to maximize and minimize the success of the immune response. Moreover, it neglected many other important parameters.

### Control and optimization problem

Building upon this work, we now seek to explore a fuller set of design parameters, over a 6-dimensional design space:

- **immune cell apoptosis rate *r*_apoptosis_:** This is the rate at which immune agents undergo apoptosis. Decreasing this could model immunoengineering to decrease immune exhaustion. (d1 in the EMEWS investigations and in Table 2)
- **oncoprotein threshold *p*_threshold_:** immune agents ignore cells with *p* < *p*_threshold_. Modulating this parameter is analogous to engineering immune cell sensitivity. (d2 in Table 2)
- **immune cell kill rate *r*_kill_:** The rate at which an adhered immune cell can trigger apoptosis in the attached tumour cell, with probability (in any time interval [t,t+Δt]) given by *r*_kill_ ·*p*·Δ*t*. (d3 in Table 2)
- **immune cell attachment rate *r*_attach_:** As described above, the rate at which an immune cell can adhere to a cell in close contact. (d4 in Table 2)
- **immune cell attachment lifetime *T*_attach_:** As described above, the mean time an immune cell maintains an attachment before searching for another target. (d5 in Table 2)
- **immune cell migration bias *b*:** As described above, this governs the randomness of immune cell chemotactic migration. This could potentially be tuned by altering chemoreceptor expression. (d6 in Table 2)

**Table 1:** Model exploration methods used with the PhysiCell/EMEWS workflows and their outcomes.

| Model Exploration Method | Outcomes in PhysiCell/EMEWS workflows |
| --- | --- |
| Active Learning (AL) | Surrogate models for characterizing the parameter space structure based on different viability thresholds |
| Genetic Algorithm (GA) | Optimal points producing the lowest final mean tumour cell counts |

**Table 2.** Workflow parameter space dimensions

| Parameter (id) [dimensions] | Min | Max | Increment |
| --- | --- | --- | --- |
| immune cell apoptosis rate (d1) [1/min] | 6.94e-6 | 6.94e-4 | 8.5882e-05 |
| oncoprotein threshold (d2) [dimensionless] | 0.1 | 1.0 | 0.1125 |
| immune cell kill rate (d3) [1/min] | 0.1 | 1.0 | 0.1125 |
| immune cell attachment rate (d4) [1/min] | 0.01 | 1.0 | 0.12375 |
| immune cell lifetime (d5) [min] | 10.0 | 90.0 | 10.0 |
| immune cell migration bias (d6) [dimensionless] | 0.1 | 0.9 | 0.1 |

We now consider two related sets of problems:

- **Cancer control:** Can we divide the 6-dimensional design space into “viable” and “non-viable” regions, where the viable region is defined to be all parameter sets (designs) for which the final cancer cell population *N*_final_ (after 21 days) did not exceed the initial population *N*_initial_, computed over multiple stochastic replicates. (**stable scenario:** *N*_final_ ≤ *N*_initial_)
- **Cancer regression:** Can we find regions of the design space where the cancer population is reduced to 10% of its initial size? (**10% scenario:** *N*_final_ ≤ 0.1 *N*_initial_) Can we find regions of the design space where the cancer population is reduced to 1% of its initial size? (**1% scenario:** *N*_final_ ≤ 0.01 *N*_initial_) Can we minimize *N*_final_? (**minimized scenario**)

To reduce the computational cost of our investigation, we explored the model in 2D; future investigations will explore optimal designs in the full 3-D model. To simulate in 2D, we only require one change: the number of immune cell agents introduced at the start of therapy (t = 14 days). We set the number of immune agents at 125: approximately the number of immune cells initially in the z = 0 plane in the full 3D model in Ghaffarizadeh et al.^15^.

We note that the edges of the hypercube represent biological and clinical constraints: biological processes (e.g., cell attachment rates) may be impossible to accelerate beyond a physical limit, while side effects may impose other clinical constraints on parameter values (e.g., if immune cells are too sensitive, they may kill healthy cells as well).

### Model exploration workflow solution: EMEWS

As previously noted, the parameter spaces of complex ABMs, coupled with the highly non-linear relationship between ABM input parameters and model outputs, require heuristic ME approaches that adaptively evaluate large numbers of simulations. Here we give a brief overview of how the Extreme-scale Model Exploration with Swift (EMEWS) framework^31^ enables the creation of HPC workflows for implementing large-scale ME studies (see^37^ for further details).

EMEWS is built on the general-purpose parallel scripting language Swift/T^38^, which provides the capability of running multi-language tasks on anywhere from desktops to peta-scale plus computing resources with a data-flow paradigm for inter-task dependencies. Central to data-flow is the run-to-completion pattern, where the outputs from completed tasks are used as inputs to subsequent tasks. While this approach for inter-task dependencies is sufficient for many applications, EMEWS introduced the ability to define resident, or stateful, tasks to encapsulate the logic within iterating, state-preserving algorithms, such as those used for ME^39^. These resident tasks can communicate with the rest of the workflow and, in fact, allow for an Inversion of Control (IoC) scheme, where the resident task can control the logical flow of the workflow, rather than needing to implement the logic in the workflow language itself. As such, an ME algorithm can be expressed in Python or R with the only required modification being that it uses the high-level queue-like interfaces, currently with two implementations EQ/Py and EQ/R (EMEWS Queues for Python and R). The queue interfaces are used for iteratively communicating parameter combinations generated by the ME algorithm to the underlying workflow for concurrent model instance execution, and for retrieving the results from those model runs. The rest of the ME algorithm remains unmodified. This allows the direct use of the many libraries relevant to ME that are being actively developed and implemented as free and open source Python and R software. We exploit this capability in the current work by implementing a Python-based GA workflow and an R-based AL workflow to explore our PhysiCell model.

The multi-language capabilities of Swift/T provide the ability to run external applications (with run to completion semantics) through the shell, in-memory libraries accessed directly by Swift/T, or Python, R, Julia, C, C++, Fortran, Tcl and JVM language applications. In this work we use the shell-based approach for launching the individual PhysiCell model runs.

## Methods: Description of computational experiments

### Description of PhysiCell and EMEWS integration

Our focus here is on delineating the shape of the parameter space with respect to the final tumour cell count. To this end, we have created two workflows: one in which the parameters to evaluate are produced by an AL algorithm^27^ and one, as a consistency check on the AL results, in which the parameters are produced by a GA ^28^. In the AL case, we employ a binary classification of regions within our space, where we define an objective and find parameter subspaces that are capable of meeting the objective (viable), versus those that cannot (non-viable). More specifically, a viable subspace contains parameters that produce final mean tumour cell counts at or below a specified threshold, and a non-viable subspace contains parameters that produce final mean tumour cell counts above that threshold. The AL algorithm trains a surrogate model (in the present case a random forest^40^ classifier) such that the viability of unevaluated regions of the parameter space can be determined without the need for additional model runs and the overall structure of the parameter space can be estimated. In contrast, the GA’s focus is on finding optimal parameters, that is, those that produce the lowest final mean tumour cell counts. While the GA is good at finding optima in complex spaces and therefore provides a useful consistency check of the AL results, unlike the surrogate model produced by the AL algorithm, it does not provide an estimate for the neighbourhood, and hence the robustness, of the solutions that it finds. Table 1 provides a summary of the two model exploration approaches we use and the types of outcomes they produce. The modularity of EMEWS allows us to easily replace one model exploration approach with the other while not affecting the rest of the workflow.

In both AL and GA cases, we run 20 stochastic variations of each parameter set, varying the random seed across the runs, and take the mean final tumour cell count over those 20 as the value to evaluate with respect to the objective threshold. Through earlier investigations we found that 20 stochastic variations provided a balance between computational effort and the stability of the evaluated outcomes‡.

In the AL algorithm, implemented in R, our goal is to iteratively pick points (i.e., parameter sets) to sample, such that we can *exploit* the results of previous evaluations, but balance that with an *exploratory* component in order to investigate undersampled regions. For the former, we use an uncertainty sampling strategy where we fit a random forest^40^ classifier on previously evaluated points, and then choose subsequent samples close to the classification boundary, i.e., where the uncertainty between classes is maximal. We cluster these candidate points of maximal uncertainty, and then select from within the clusters, in order to ensure a level of diversity in the sampled points and, therefore, a greater expected reduction of uncertainty^41^. The exploratory component randomly samples points in the parameter space. At each iteration of the AL algorithm, all of the chosen points, i.e., exploit and explore points, are collected and evaluated in parallel. The results of those evaluations are gathered and the random forest model is refit with the additional data, allowing for a new set of points to be chosen for sampling. This process is continued until a convergence or, in the present case, a maximum number of iterations is achieved.

Random forest classifiers can also produce measures of relative importance of a space’s dimensions. A random forest classifier is an ensemble of decision trees, where each tree is trained on a subset of the data and votes on the classification of each observation variable. Importance can be calculated from the characteristics of decision trees. A commonly used measure for importance is the mean decrease in Gini, a measure of the weighted average of a variable’s total decrease in node impurity (which translates into a particular predictor variable’s role in partitioning the data into the defined classes). Gini decrease is an effective measure of the relative importance of a variable in classifying the target observation, across all of the decision trees in the forest. A higher Gini decrease value indicates higher variable importance and vice versa and we have ordered the dimensions in our output plots accordingly (see Results, Figures 2–4). Details of the AL algorithm are further described in37 and the R code can be found in GitHub; see the link in the Electronic Supplementary Material.

**Figure 1:**
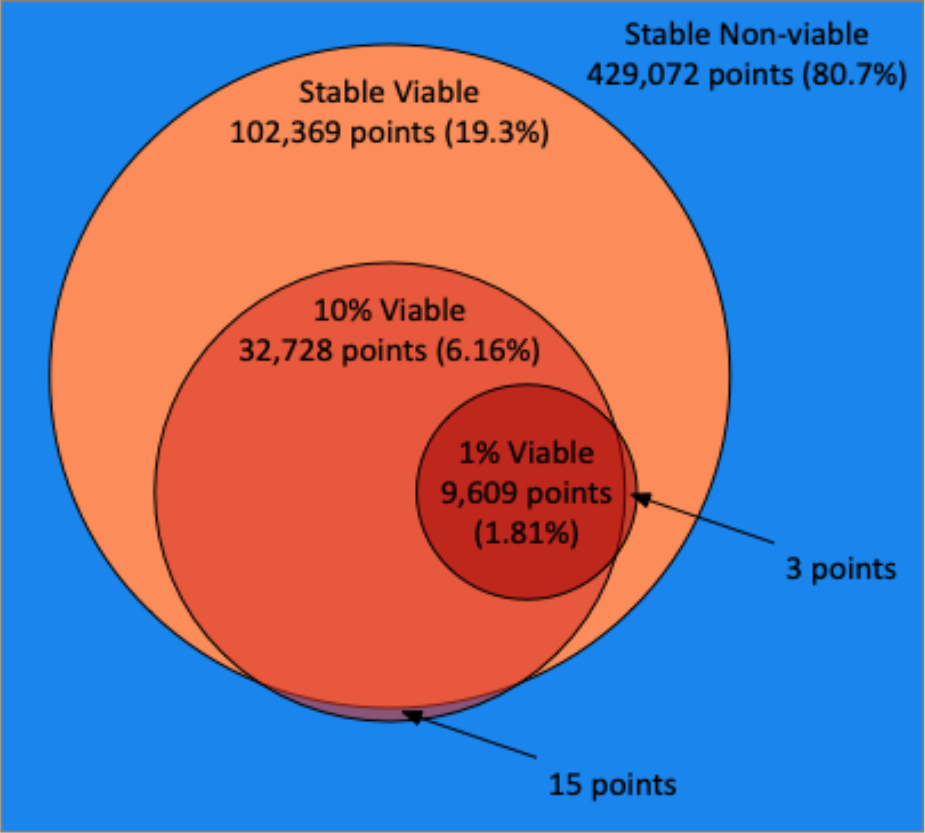
Venn diagram illustrating the total points, proportions, overlaps and disagreements between the classifications of the PhysiCell model parameter space made by the random forest models trained via the three AL scenarios (stable, 10%, 1%).

**Figure 2:**
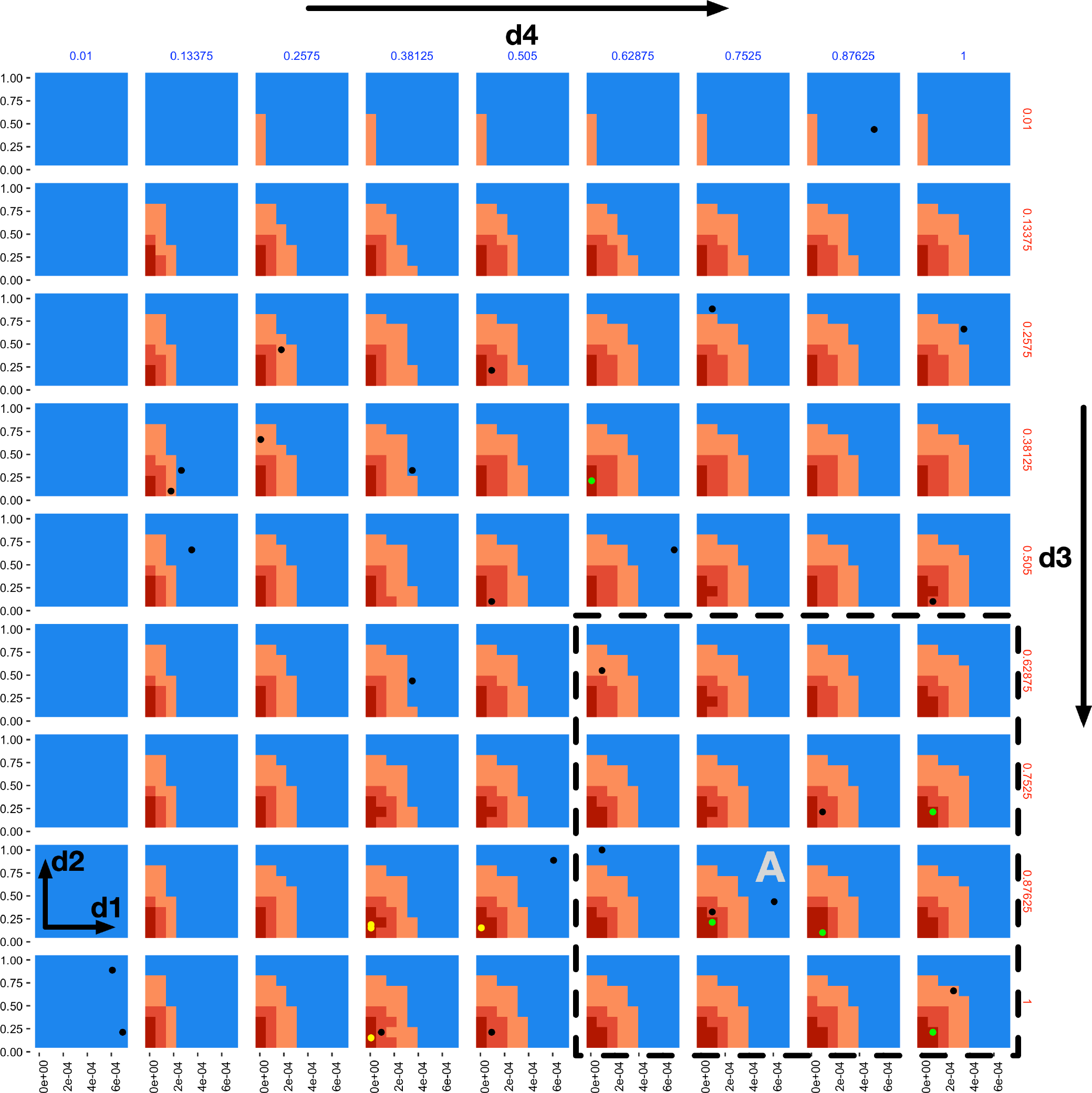
A 4-dimensional slice of the 6-dimensional parameter space after 20 iterations of all three AL scenarios. Immune apoptosis rate (d1) and oncoprotein threshold (d2) are plotted against each other in each of the individual subplots. The immune kill rate (d3) is varied along the subplot grid rows and the immune attachment rate (d4) along the subplot grid columns. The immune attachment lifetime (d5) and immune migration bias (d6) parameters are set at d5 = 80 minutes and d6 = 0.8. The space is coloured based on the classifications of the three AL scenario random forest models, where the blue regions were classified as non-viable by the stable scenario model (the tumour population grew), the light orange regions were classified as viable by the stable scenario model (the tumour population did not increase), the darker orange regions were classified as viable by the 10% scenario model (the final tumour population was under 10% of the initial value) and the dark red regions were classified as viable by the 1% scenario model (final tumour population under 1% of the starting value). The green and black dots indicate points where simulations were run, resulting in viable and non-viable outcomes for the 1% scenario, respectively. The yellow dots correspond to final GA population points from the seeded GA scenario.

**Figure 3:**
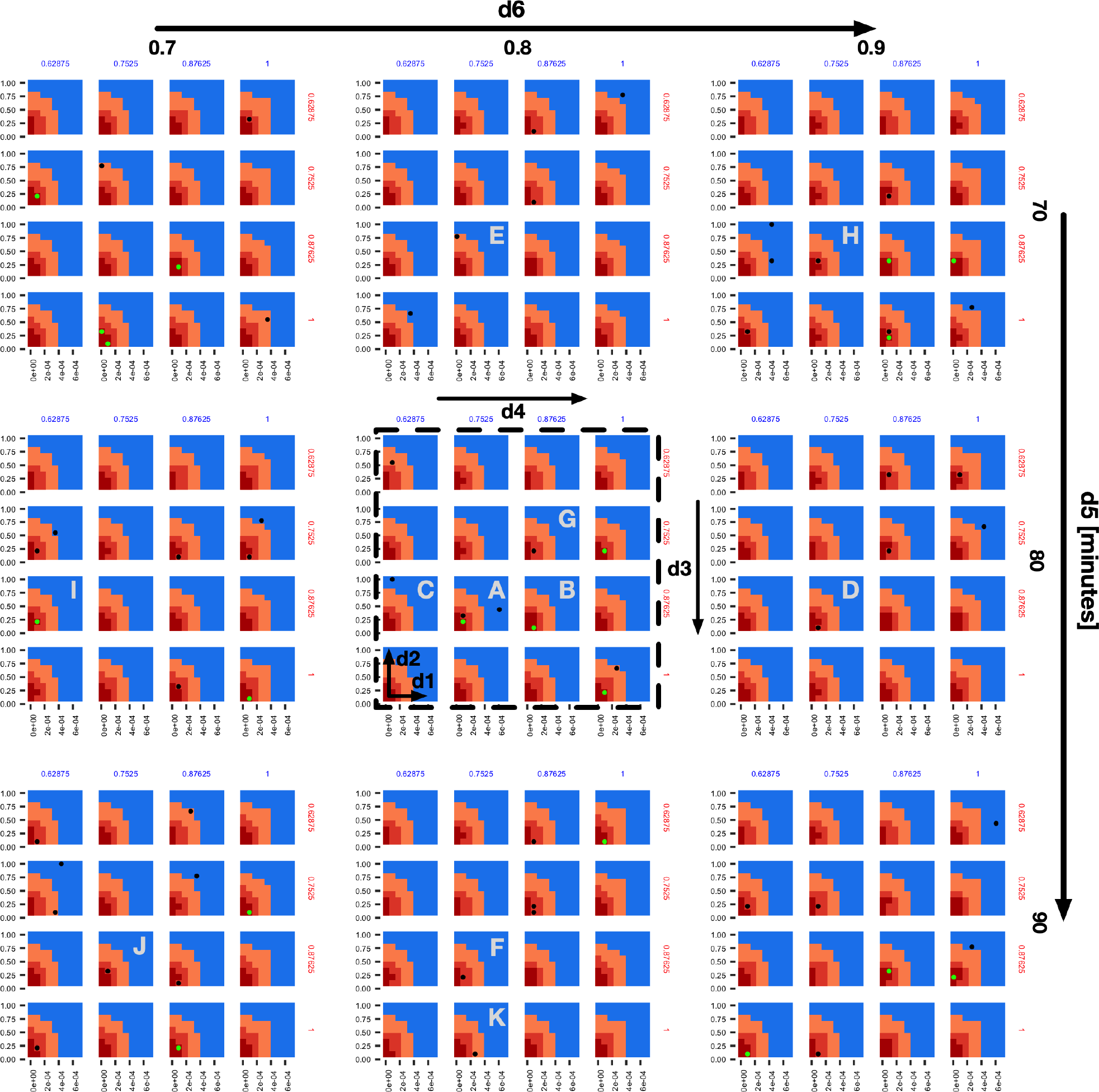
An expansion of the 4×4 subplots demarcated with a dashed bounding box in the bottom right of Figure 2, here also shown with a dashed bounding box, to include variations along the d5 (overall rows) and d6 (overall columns) dimensions. The space and dot colours are defined as in Figure 2. Subplots B-F are those that contain sampled points and are Manhattan distance 1 neighbours of subplot A, and subplots G-K are the ame but at Manhattan distance 2.

**Figure 4:**
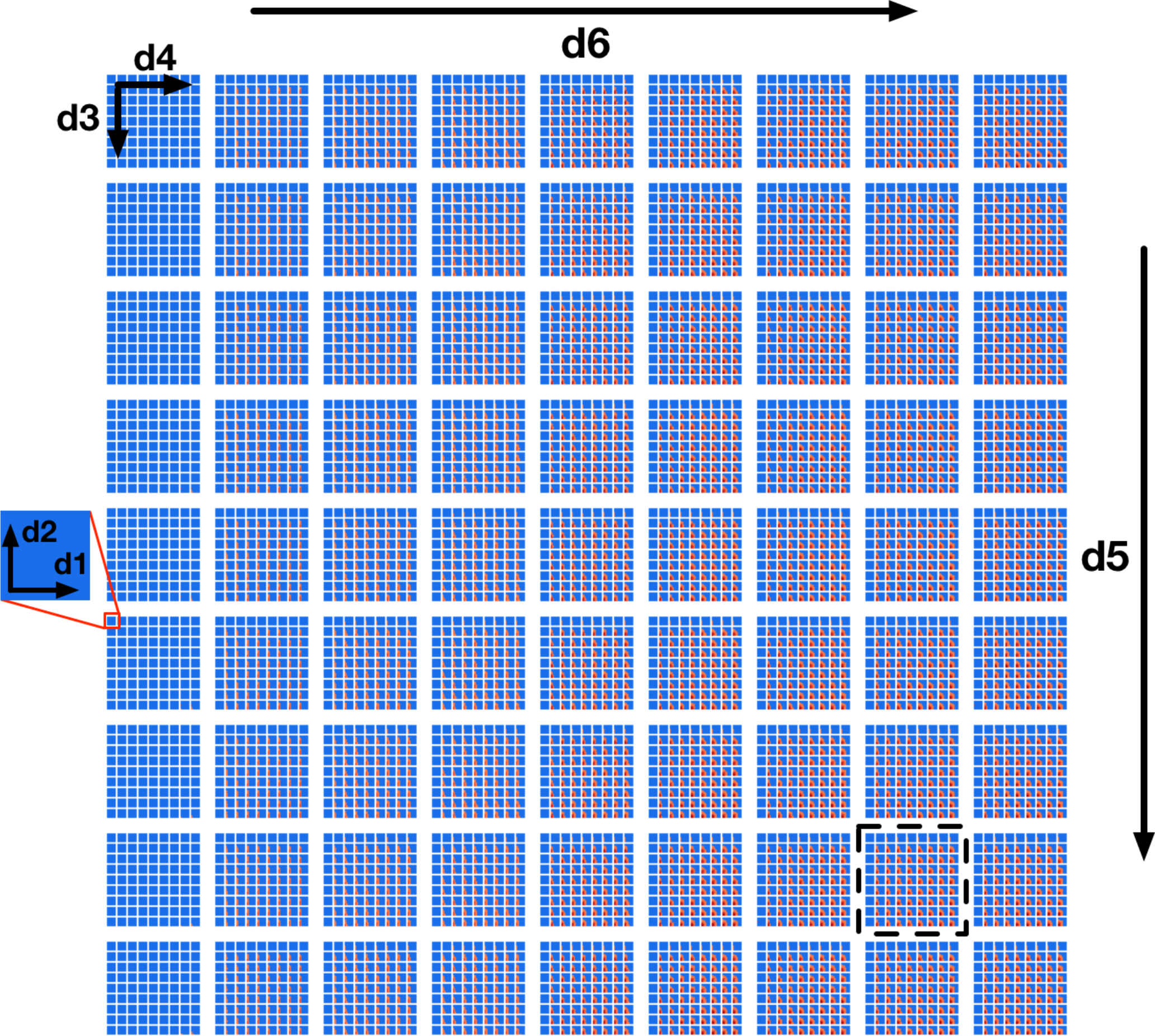
**A full 6-dimensional view of the design space**, where the space colours are defined as in Figure 2 and the location of the slice of the space shown by Figure 2 is illustrated by a dashed bounding box. d5 and d6 vary along overall rows and columns, and d3 and d4 vary along the sub-grid rows and columns, and d1 and d2 vary along x and y axes within the individual sub-grids.

In our GA algorithm, our goal is to iteratively evolve a population of points to produce points that when evaluated yield the lowest mean final tumour cell counts. We have implemented this using the DEAP [REF] evolutionary computation Python framework, using the evolutionary algorithm presented in chapter 7 of^42^. During each iteration, the best points from the currently evaluated population are selected using a tournament selection method to create a new population. Each of these points is then mated with another according to a crossover probability and, finally, each of the resulting points is mutated according to a mutation probability. At each GA algorithm iteration, the new population is evaluated and the evaluation results are gathered. Here, as with the AL case, we make use of the ability to concurrently evaluate as many design points as the HPC resource allocation provides. Also similarly to the AL case, the workflow continues until the desired number of iterations are achieved.

Both the AL and the GA ME algorithms are integrated into EMEWS workflows using EMEWS queues, EQ/R and EQ/Py, respectively. The points to evaluate for each iteration are passed from the ME algorithms to the Swift/T workflow for evaluation via the queues. As mentioned earlier, each of these parameter points is expanded into 20 stochastic variations and those variations are all run in parallel. When each individual run completes, the workflow executes a small script (written in Python) to parse the final tumour cell count from the PhysiCell simulation output file. The tumour cell counts for each of the 20 variations are collected and the mean is calculated using another small script (written in R)§. This mean tumour cell count is then passed back to the ME algorithm as the result of the evaluation, again via the queues.

The PhysiCell simulation itself is a stand-alone command line application that takes the path to an XML format file, containing all the parameter input for a simulation run, as an argument. The workflow launches the PhysiCell application using a shell script, passing the path to a specially constructed XML parameter file. This XML file is created for each run by reading a base XML file that contains a default set of parameters, replacing only the parameters of interest with those produced by the ME, and writing the new XML to a location where it can be read by the PhysiCell application. This transformation is implemented in a small amount of Python code, executed by the workflow prior to launching a simulation run.

Using the two EMEWS workflows, we performed the following experiments. For the AL case, we ran 3 scenarios using 3 classification thresholds. In the first scenario, viable subspaces were those that produced a stable tumour cell count, i.e., non-progression of the tumour (*stable scenario*). Since the initialized tumour size consists of 900 tumour cells, the AL classified all runs where the final mean tumour cell count was less than our simulation starting value of 900. The second scenario examined a case where the immunotherapy resulted in a reduction of the initial tumour to 10% of its size; this is the *10%* scenario which classified those runs with < 90 live tumour cells. The third scenario examined situations where immunotherapy reduced the tumour to 1% of its original size; this is the *1% scenario* which captured runs with < 9 live tumour cells at their conclusion. For each scenario, the AL algorithm was run for 20 iterations, sampling 25 points using the maximal uncertainty strategy and 25 using the random strategy, as described above. We performed 20 simulations for each point, varying the random seed. The full parameter space from which we sampled consisted of all the combinations of our 6 parameters, with each parameter discretized across 9 different values (Table 2). Each AL algorithm was seeded with an initial design, a pre-evaluated set of points. The stable scenario was seeded with 707 evaluations whose points were sampled using a Latin Hypercube Sampling (LHS) strategy. The 10% and 1% scenarios were seeded with the 1000 evaluations (i.e., 50×20 iterations) performed by the first scenario.

For the GA case, we ran two scenarios differentiated by the composition of the initial population. In the “seeded” scenario, the initial population consisted of 12 unique points found by the first AL scenario (i.e., stable scenario) to be the best (i.e., lowest mean tumour cell count) and 38 randomly selected points. The second “unseeded” scenario consisted of only randomly selected initial points. The space from which points are selected is defined by the minimum and maximum values of our 6 parameters (see Table 2). Both scenarios had population sizes of 50 and were run for 30 iterations. The mutation probability was set to 0.2 and the crossover probability to 0.5 for both scenarios.

## Results

All experiment scenarios presented in the previous section were performed on the Cray XE6 *Beagle* at the University of Chicago, hosted at the Argonne National Laboratory. *Beagle* has 728 nodes, each with two AMD Opteron 6300 processors, each having 16 cores, for a total of 32 cores per node; the system thus has 23,296 cores in all. Each node has 64 GB of RAM. The workflows were run over 126 nodes, using 8 processes per node. 8 PhysiCell simulations were concurrently run on each node with 4 threads allocated to each simulation, thus utilizing all 32 cores on a node. With a total of 126 nodes, we were able to run a maximum of 1005 (126 nodes * 8 simulations per node - 3 processes for workflow and ME overhead) simulations in parallel.

In the AL stable scenario, 1707 points in total were evaluated (including the 707 seeded points), with 394 points (23.1%) as viable and 1313 as non-viable. Out of the 531,441 total points that made up the discretized parameter space, the final random forest model classified 102,369 points (19.3%) as viable. In the 10% scenario, 2000 points in total were evaluated (including 1000 seeded points), 297 (14.9%) as viable, and 1703 as non-viable. Out of the 531,441 total points, the random forest model classified 32,728 points (6.16%) as viable. We note that 15 of the points classified as viable in the 10% scenario were classified as non-viable in the stable scenario, which points to the imperfection of the random forest models. Indeed, one should not expect that a surrogate model would replicate with full fidelity the more complex stochastic simulation that is being approximated. Nevertheless, this represents a very small misalignment when considering the size of the full discretized space. In the 1% scenario, 2000 points in total were evaluated (including 1000 seeded points), 204 as viable (10.2%), and 1793 as non-viable. Out of the 531,441 total points, the random forest model classified 9,609 points (1.81%) as viable. Here all of the viable points were also classified as viable by the stable scenario, although 3 of the points were classified as non-viable by the 10% scenario. A Venn diagram illustrating the relationships between the space classifications made by the three AL scenarios, including their overlaps and disagreements is shown in **Error! Reference source not found**‥ The three AL scenarios each took between 10-12 hours to run, accounting for 40k-48k core-hours each, clarifying our need for employing an HPC workflow approach.

A “slice” of the 6-dimensional parameter space after 20 iterations of all three AL scenarios is shown in Figure 2 as a grid of 2D plots. This visualizes regions of the parameter space meeting the classification criteria in a fashion that incorporates the relative importance of each parameter based on the dimension’s Gini decrease value. The two most important dimensions, immune apoptosis rate (d1) and oncoprotein threshold (d2) are plotted against each other in each of the individual subplots, which are laid out in a grid. The next two important parameters, immune kill rate (d3) and immune attachment rate (d4), are the axes along which the grid of subplots are plotted; the immune kill rate for the subplot grid rows and the immune attachment rate for the subplot grid columns. The final two, least important dimensions, immune attachment lifetime (d5), and immune migration bias (d6) are kept constant at d5 = 80 minutes and d6 = 0.8.

The space is partitioned into different regions based on the evaluation results of the 3 different random forest models trained on the 3 different AL scenario data sets. The blue regions were classified as non-viable by the stable scenario model. The light orange colour marks all the points (both evaluated and not) classified as viable by the stable scenario model. The darker orange marks points classified as viable by the 10% scenario model and the dark red are those classified as viable by the 1% scenario model. One can observe the steeper changes of the parameter space characterizations within each subplot in the d1 direction when compared to d2, and slow changes along the d3 and d4 dimensions, providing support to the dimensional importance ordering we are using. Individual parameter sets evaluated in the 1% scenario can be seen as green (viable) and black (non-viable) dots.

In the subplot labelled “A” (Figure 2) we can see how the AL algorithm is attempting to characterize the boundary of the 1% scenario space. The green and black dots along the boundary are points produced by the AL algorithm as it tries to resolve regions of maximum uncertainty and, in doing so, delineate the boundary. The remaining black dot in the blue region of the subplot is an example of the AL’s random selection explore strategy. To provide a better sense of boundary finding in all 6 dimensions, in Figure 3 we expand our perspective to include the 4×4 subplots demarcated with a dashed bounding box in the bottom right of Figure 2, along with their 8 neighbours in the d5 and d6 dimensions. Here, d5 is varied along the overall rows, with values of 70, 80 and 90 minutes, and d6 is varied along the overall columns, with values of 0.7, 0.8 and 0.9. The original demarcated area from Figure 2 is placed at the centre. Figure 3 shows neighbours of our original plot A (from Figure 2), where we have identified subplots with sampled points that are distance 1 neighbours (in Manhattan distance) with B-F and those that are distance 2 neighbours with G-K. While not visible in Figure 2, we observe that points are evaluated along the classification boundary across the d5 and d6 dimensions as well.

Lastly, Figure 4 provides a zoomed-out view over the entire space (full 6-dimensional space) characterized by the random forest classifiers where, instead of fixing the d5 and d6 parameter values, we vary them across the overall rows (d5) and columns (d6). Each subplot in Figure 4 displays a sub-grid of 2D plots with the same axes as in Figure 2: d1 and d2 plotted against each other with d3 and d4 varying along each overall sub-grid row and column. The sub-grid corresponding to Figure 2 is highlighted by a dashed bounding box.

We note that the characterization of the parameter space in Figures 2–4 is based on the use of the 0.5 probability threshold between viable and non-viable classifications, as generated by the random forest models. The sample F-score values obtained with the 0.5 probability threshold through 10-fold cross validation at the end of the stable, 10%, and 1% scenarios were 0.87±0.04, 0.80±0.05 and 0.64±0.08, respectively. Adjusting the classification threshold value, such that a higher threshold would be required for classifying a point as viable, could yield higher positive predictive values at the expense of increased false negatives, but may be worth considering from the point of view of the increased confidence within the viable regions.

Our GA workflow was run as a consistency check against the AL workflow results. Moreover, GA represents a more traditional approach to this type of optimization problem, when lacking the AL approach and resources to characterize the full parameter space. In the first scenario where the initial population was seeded with the 12 best results from the stable AL scenario, the final GA population, after 50 iterations, consisted of 16 unique points with an average mean tumour count of 0.764, a standard deviation of 1.112, a minimum value of 0 (the optimal solution) and a maximum value of 4.75. The unseeded scenario consisted of 41 unique points with an average mean tumour count of 1.411, a standard deviation of 2.352, a minimum of 0 and a maximum of 14.05. The final seeded population while better than the unseeded has less breadth likely due to the initial constraints on the population. In Figure 2, the yellow dots correspond to 4 points from the final seeded scenario population, and are all located within the AL 1% boundary, as would be expected. In contrast to the AL workflow, the GA, while good at finding optimal points, does not help in delineating the parameter space nor provide an estimate of the robustness of the produced solutions. Furthermore, it does not generate a model, such as the AL’s random forest classifier, that can be used to classify points without running any additional simulations.

## Discussion and Future Directions

This work extended our prior proof-of-concept implementation^16^ to demonstrate the utility of integrating the PhysiCell and EMEWS frameworks to iteratively explore a high-dimensional therapeutic design space and optimize a complex cancer immunotherapy model. In particular, we were able to investigate highly relevant clinical problems in cancer immunotherapy: given a therapeutic design space with biological and clinical constraints, can we identify optimal designs to minimize the remaining number of tumour cells, can we characterize the robustness of those optimal designs, and can we determine the most important design parameters?

As in more traditional approaches, we applied GAs to find the treatment optima within the space. Interestingly, there were multiple parameter sets (therapeutic designs) that were able to completely eliminate the live cancer cells by the end of the simulated treatment, which matches clinical observations that immunotherapies can lead to complete responses in some patients^2,43,44^.

We found that these optimal designs were on and near the boundary of the design space hypercube. When optima are found within the interior of a design space, it indicates that finding the ideal balance between design parameters is most important to improving the design. When optima are found on the boundary of a constrained design space rather than the interior, it indicates that the optima could be improved by relaxing the constraints on one or more design parameters. In the case of the immunotherapy design problem, these constraints are primarily biological (e.g., biological limits on immune cell lifetimes and killing rates) and clinical (e.g., limits imposed by toxicity). This suggests that improving these constraints—e.g., reducing toxicity by improving the specificity of immune-cell targeting—may be key to improving therapeutic response in more patients.

While finding optima on the constraint boundaries may have been expected due to the lack of explicit negative model feedbacks (e.g., *de novo* cancer cell mutations), the result was by no means certain for the highly nonlinear model. For instance, Ozik et al.^16^ found that the migration bias (d6) had a nonmonotonic influence on the treatment success in the full 3-D model: large and small values of d6 (corresponding to highly exploitative or highly exploratory immune cell migration) gave improved responses over more mixed exploitation-exploration migration strategies.

Even with knowledge of the individual design constraints, it is the *interaction* between the parameters that drives the success or failure of any given design. While the optima were near the boundaries of most of the dimensions in the design subspace, they were in the interior of d4. Hence, the balance between d4 and the other parameters was important in achieving these optima. It would have been difficult to anticipate the specific interactions between the parameters and constraints based upon single-parameter data *a priori*.

Moreover, the integrated PhysiCell-EMEWS framework—combined with HPC resources—enabled previously infeasible investigations. By using AL to guide our sampling of the design space, we were able to move beyond finding optima to understand the *topology* of the design space by characterizing increasingly aggressive treatment goals. The stable design space—where the cancer was kept in control—included just 20% of our initial constrained design space. Increasing the goal to eliminating 90% of cancer cells drastically shrank the viable therapeutic designs to 6% of the space. The decreasing marginal utility can be seen when moving to the goal of eliminating 99% of cancer cells: the viable design space is now under 2% of the original tested design space. This characterization of the therapeutic design space would not have been possible without using active learning to guide the sampling of parameter space, even on HPC resources. Our model findings are qualitatively consistent with the performance of clinical trials, where far fewer patients experience a complete response than those whose cancers are controlled by immunotherapies. For example, Carretero-González et al. reported that in a meta-analysis of twelve Anti-PD1/PD-L1 trials, only 2.19% patients achieved a complete response, while an additional 44.56% of patients achieved partial response or stable disease.^44^

We note that this approach supports a bootstrapping investigation: identifying the viable control designs can seed the search for the 90% cell kill regime, which in turn can seed the search for more aggressive treatment goals. The distance between the edges of these identified treatment subspaces helps to characterize the robustness of the designs.

The optima (identified by the GA) lie within the extremely small 99% cell kill regime, showing that they may not be particularly robust to engineering variability and evolution of the cancer cells. Missing this narrow design regime could have negative long-term clinical consequences: eliminating 99% or 90% of cancer cells would place a strong selective pressure on the cancer cells, encouraging the development of therapeutic resistance. It may well be wiser to target the control case, to improve the patient survival times^6^. Such results have similarly been suggested by evolutionary game theory^45,46^.

As an additional benefit of the AL-based model exploration over GA workflows, the random forest classifier can rank the importance of the parameters in determining the success or failure of treatment designs. The most important parameter was d1: the immune cell death rate. Minimizing d1 is equivalent to maximizing the immune cell life time, and thus its maximum number of cell kills. Minimizing d1 is analogous to reducing T cell exhaustion and increasing T cell killing capacity, one of the most active areas of research in cancer immunology^47–49^. The second most important parameter was the immune cell detection floor (d2): decreasing d2 corresponds to increasing immune cell sensitivity. Increasing immune cell recognition is also an extremely active area of work in cancer therapy. Interestingly, the optimal therapy parameters occurred along the minimum allowed values of d1 and d2, and hence the simulated treatments were limited by biological constraints (d1) and clinical constraints (d2). The minimum value of d2 could only be reduced further by increasing the specificity of immune cell response^5,50^. It is also interesting that varying d5 (tumour-immune attachment time) was identified as having little impact on the treatment success. Interestingly, discussion with cancer immunologists has suggested that tumour-T cell contact times are relatively brief. However, detailed experiments have found both fast (< 200 min) and slow (> 200 min) contact times for cell kills^51^, while other work suggests that extended contact times can improve cytotoxic responses^52^. It is likely that if we refined our constraints on *r*_kill_ (d3) to reflect such detailed measurements, longer contact times (d6) would be required to achieve effective responses.

It is striking that the model identified rankings of the important and unimportant design parameters based solely on a model of physical interactions, without explicitly modelling the molecular biology of the system. This validates the approach of using iterative, high-throughput simulation investigations of physics-inspired models to understand and optimize the behavioural rules. Once optimal rules are identified, the focus can turn to identifying molecular mechanisms that can be linked to the cell behaviours.

For instance, the immune cell lifetime (corresponding to 1/d1), immune cell sensitivity to detecting an adhered cell as immunogenic (d2), and the immune cell killing rate (d3) were the three most important design parameters. Future agent-based models could include more detailed models of immune cell signalling, including stimulatory pathways (e.g., receptors that prime immune cells) and inhibitory pathways (e.g., receptors that suppress cytotoxicity, suppress cycling, increase apoptosis, or otherwise contribute to exhaustion). Mathematical models of receptor pathways have found that the sensitivity, rate of activation, and duration of response depend upon key rate parameters for receptor-ligand binding, dimerization, internalization, turnover, synthesis, and decay (e.g., see references^53–55^). With detailed models of immune receptor dynamics, we could tune and balance these parameters as determined by the earlier ABM investigations, and hence “implement” the rules for d1, d2, and d3. Future models could incorporate such models in each individual immune cell agent to engineer their activation across space of the tumour, and perhaps even to maximize their activation near tumours, rather than in adjacent tissues where immune cell activation could contribute to immunotherapy toxicity.

There are limitations in the current work. First, while the abstract “immune cell” PhysiCell agents allowed us to focus on the physical limits of tumour-immune contact interactions, we must improve the biofidelity by explicitly modelling key immune cell types, particularly T cells and dendritic cells. This would facilitate more direct comparisons with known cancer immunology. Molecular-scale biology, such as immune receptor binding dynamics, should be incorporated into the individual cell agents to better incorporate current molecular-scale hypotheses on immune cell function and tumour cell recognition. Moreover, this investigation did not include interactions with the stroma, particularly fibroblasts and matrix remodelling, the vasculature and angiogenesis, and inflammatory processes^56–58^. These should be included to better understand the dynamics of cancer-immune interactions in 3-D environments that more closely resemble primary and metastatic tumours in patients. These higher-fidelity models could be tailored to specific clinical trials and experiments, allowing more direct, quantitative comparison between simulation predictions and clinically observed tumour-immune interactions.

Rather than starting with an initial distribution of cancer phenotypes, models should account for continuous genetic and epigenetic variability in cancer cells that can drive treatment failure and unexpected tumour-stroma interactions. Relatedly, future models should include normal tissue components to better model adverse effects of highly cytotoxic immune therapies; these would shift the optimization landscape.

Lastly, we know that there are artefacts associated using 2-D simulations to investigate 3-D problems; these artefacts could produce misleading rankings of the influence of various parameters. For example, the randomness of migration (d6) may have more benefit in exploring space to find tumour cells in 3-D than in 2-D. However, high-throughput 2-D simulation investigations as in this paper could be used to help focus subsequent 3-D investigations.

The current study was limited to 2D rather than 3D in part because PhysiCell—as most current ABM platforms for complex multicellular problems—is optimized for fast computation on shared memory architectures such as desktop workstations or single HPC nodes (via OpenMP^59^), whereas most HPC platforms are optimized for distributed memory applications. Thus, individual model instances cannot currently be accelerated by extending them onto multiple HPC nodes (e.g., via MPI^60^). Moreover, most biological ABMs integrate molecular-scale models using standards such as the systems biology markup language (SBML), and by coupling with SBML solvers such as *libRoadrunner*^61^. However, these ABM+SBML models have generally not been tested on large-scale HPC platforms.

Future ABMs will need to be redesigned for HPC architectures, such as combining HPC-tailored ABM engines (e.g., RepastHPC^62^) with the APIs and syntax of existing biology-focused ABMs. This is especially needed for future ABMs that will model not just multiple immune cell types, but also additional dynamics in the microenvironment and (receptor) signalling dynamics in each cell agent, which will require simulating tens of nonlinear ordinary differential equations at small time steps.

Ideally, ABMs will be co-designed with upcoming exascale platforms, to not just incorporate hybrid OpenMP-MPI architectures (to efficiently parallelize single simulations over multiple HPC nodes), but to also leverage the unique memory architectures and onboard GPU and other accelerators that could facilitate the molecular-scale detail in these models^63–65^. Co-design focused on challenging cancer immunology problems could drive cutting-edge technological advances in multi-scale, multi-physics, and discrete-continuum modelling software and benchmark exascale computing platforms.

Now that we have fully demonstrated the learning-accelerated PhysiCell-EMEWS platform, we will turn our attention to improving the model. In particular, we will work closely with cancer immunologists to refine the immune cell models and more explicitly model T cells, dendritic cells, and other immune players in cancer immunotherapy, as discussed above. We will also investigate the role of continuing tumour variability in driving T cell exhaustion and related immune escape processes.

We plan further refinements to the model exploration pipeline as well. The increased model detail will come at the cost of even higher-dimensional parameter and design spaces. We will explore the possibility of using the simulation runs to build surrogate models that map from the model parameters to the simulation metrics (thus far total live cell population, but potentially including entropy measures of tumour heterogeneity^66,67^, emerging measures of the tumour ecology^68^, and measures of impact on nearby non-tumour cells). We could use apply dimensionality reduction techniques^69^ to the surrogate model to eliminate redundant parameters and refine our investigation of the original model. These surrogate models could also incorporate heteroskedastic stochastic variance^70^. We will also explore extensions of PhysiCell to HPC to enable higher-fidelity simulations that more closely mirror real biological systems. Such higher-fidelity models could be explored in high throughput (and potentially in 3D) on current and emerging Top500 HPC systems.

## Conclusions

The use of simulation is an integral component of the modern engineering workflow. While the ability to determine the sufficient level of fidelity for simulations of biological processes is still an open area of investigation, we believe that developing simulation-based methods for potentially engineering mechanism-based biomedical interventions should not wait until the former situation is “solved.” In fact, we contend that investigating the of modes and methods of simulation-aided biomechanistic engineering can aid in the determining of how detailed biosimulations “need” to be. Even at the current level of abstraction, the ME and optimization examination presented herein identifies non-intuitive insights that may help guide concurrent work on improving one of the most currently promising emerging cancer therapies.

## Acknowledgements

PM and RH were funded in part by the National Science Foundation (Award 1720625), the National Institutes of Health (U01-CA232137-01), the Breast Cancer Research Foundation, and the Jayne Koskinas Ted Giovanis Foundation for Health and Policy. JO, NC and GA were funded in part by the National Institutes of Health (R01GM115839 and R01GM121600). This work was completed in part with resources provided by the Laboratory Computing Resource Center at Argonne National Laboratory (the Bebop cluster), and the University of Chicago (the Beagle supercomputer).

## Notes

† A free nanoHUB registration is required to run the model.

‡ While not within the scope of the current study, future work will be able to examine the impacts of the number of stochastic variations on the estimated shape of the parameter space boundaries, where variable stochastic replicates could be used to account for heteroskedasticity.

§ Here we note the capabilities that Swift/T provides in being able to integrate multi-language components for different aspects of a workflow. We utilized Python and R for the different scripts purely for convenience.

